# Factorial variation of saccade vigor with dual decision processes

**DOI:** 10.1101/2025.08.25.671783

**Authors:** Wanyi Lyu, Harry Parmar, Thomas R. Reppert, Jeffrey D. Schall

## Abstract

Canonical stochastic models of decision-making treats decision and action as independent and sequential processes. However, studies involving limb movements consistently show that movement duration and kinematics are influenced by the quality of evidence. We tested whether saccade velocity varies with the quality of evidence in monkeys performing a visual search GO/NOGO task in which singleton elongation cued the GO/NOGO stimulus-response rule and the location of a color singleton specified saccade endpoint. We factorially manipulated the efficiency of stimulus-response cue discrimination by varying the elongation of the singleton and the efficiency of singleton localization by varying the color similarity between the singleton and distractors. The effectiveness of the manipulations was revealed by the response times on correct trials that were separately modified by the singleton localizability and stimulus-response cue discriminability, by the incidence of localization and response selection errors with separately modified error response times. Saccade velocity was higher on correct relative to error trials and was inversely proportional to response time. Saccade velocity was separately modified by singleton localizability and stimulus-response cue discriminability. Distinct patterns of error rates and saccade velocity across monkeys indicated individual differences in decision-making strategies. These findings demonstrate that the process selecting endpoints can influence both the timing and dynamics of saccadic eye movements. Incorporating saccade vigor can provide valuable constraints on biologically plausible decision models and help address the persistent challenge of model mimicry.

## Introduction

Canonical sequential sampling stochastic accumulator decision models explain systematic and stochastic variation of response time (RT) in terms of the dynamics of evidence accumulation to a threshold (Forstmann et al., 2016; Ratcliff, 1978; Ratcliff et al., 2016). Recent research with manual responses has described systematic variation in execution as well as in RT (Gallivan et al., 2018; Molano-Mazón et al., 2024; Servant et al., 2015, 2021; Wispinski et al., 2020). Such variation in execution dynamics can be explained by models that incorporate dynamics of both evidence accumulation and responses execution (Dendauw et al., 2024; Molano-Mazón et al., 2024). Unlike manual movements, saccade dynamics are commonly thought to vary only with amplitude in strict association with duration. However, recent research has demonstrated systematic, if small, variation in saccade velocity with confidence or urgency (Seideman et al., 2018; Thura, 2020), motivation (Reppert et al., 2015; Shadmehr et al., 2019), speed-accuracy trade-off (Reppert, Servant, et al., 2018), and signal strength (Molano-Mazón et al., 2024). Vigor – defined as saccade peak velocity normalized for saccade amplitude – has been used to describe the strength, effort, or energy expenditure of a movement (Spering, 2022).

We tested how saccade vigor varied in a double factorial visual search task designed to separately modify two distinct evidence accumulation processes (Lowe et al., 2019). Macaque monkeys earned reward for shifting gaze to a color singleton unless the search stimuli were square. Variation in the elongation of the search stimulus defined the first evidence accumulation process, which determines the appropriate response (GO vs. NOGO). Variation in the chromatic similarity of the singleton and distractors defined the second evidence accumulation process, which determines the singleton location. Stimulus elongations and color similarities were chosen to allow high and low efficiency of both processes, creating a 2x2 factorial design.

## METHODS

### Data acquisition

This study complies with the ARVO Statement for the Use of Animals in Ophthalmic and Vision Research. All procedures were approved by the Vanderbilt Institutional Animal Care and Use Committee in accordance with the United States Department of Agriculture and Public Health Service Policy on Humane Care and Use of Laboratory Animals.

Two macaques (Macaca radiata), identified as Da (Male, 12.8 y/o at the time of study, 8.0 kg) and Le (Male, 7.5 y/o, 13.1 kg), performed a GO/NOGO visual search task. Eye position was sampled (1000 Hz) with video-oculography (Eyelink, SR Research). Monkeys earned fluid reward for shifting gaze to a color singleton presented with seven distractor items in a circular array, unless the stimuli were squares. Trials began with fixation of a central stimulus. After a minimum interval, random non-aging delay between 800-2000 ms, eight iso-eccentric, equiluminant, square or rectangular stimuli were presented at 6° or 8° eccentricity. The singleton and distractors were square on ∼25% of trials, cuing a NOGO response, and monkeys were rewarded for maintaining fixation at the central spot for 1000 ± 0 ms. If stimuli were elongated, then monkeys were rewarded for shifting gaze to the singleton and maintaining fixation for 800 ± 0 ms (Le) or 1000 ± 0 ms (Da). The inter-trial interval was fixed at 2 seconds. Singleton localization and stimulus-response cue discrimination were separately modified by manipulating visual search stimuli along two dimensions (Figure 1). Variation of singleton-distractor color similarity manipulated the efficiency of singleton localizability. To discourage monkey from establishing a target template, singleton and distractor colors were drawn randomly from four chomaticities: red (CIE x=628, y=338, Y= or x=604, y=339, Y=5.2), off-red (x=552, y=399, Y=4.5 or x=520, y=405, Y=6.6), green (x=280, y=610, Y=4.6 or x=292, y=575, Y=6.1), and off-green (x=322, y=558, Y=4.6 or x=364, y=426, Y=6.8) presented on a gray background (x=275, y=228, Y=0.54 or x=334, y=375, Y=0.6). Stimulus elongation manipulated the efficiency to discriminate the stimulus-response cue. Stimuli were randomly drawn from four aspect ratios: 1.0 for NOGO and either 1.2-1.4 (less elongated and harder) or 2.0 (more elongated and easier) for GO. Orientation was counterbalanced with vertical (horizontal) cuing GO for monkey Da (Le). The 2 manipulations x 2 levels of difficulties resulted in four types of GO trial: the easiest was high efficiency singleton *localizability* with high efficiency response-cue *discriminability* (**H_Local_H_Discrim_**). The most difficult was low localizability with low discriminability (**L_Local_L _Discrim_**). Of intermediate difficulties was low localizability with high discriminability (**L_Local_H_Discrim_**) and high localizability with low discriminability (**H_Local_L_Discrim_**). The two NOGO trial types are high efficiency singleton localizability (**H_Local_O**) and low efficiency singleton localizability (**L_Local_O**). In total, there were 10 singleton-distractor color combinations x 3 response cues: square, less elongated, and more elongated (Table 1).

**Figure 1.**
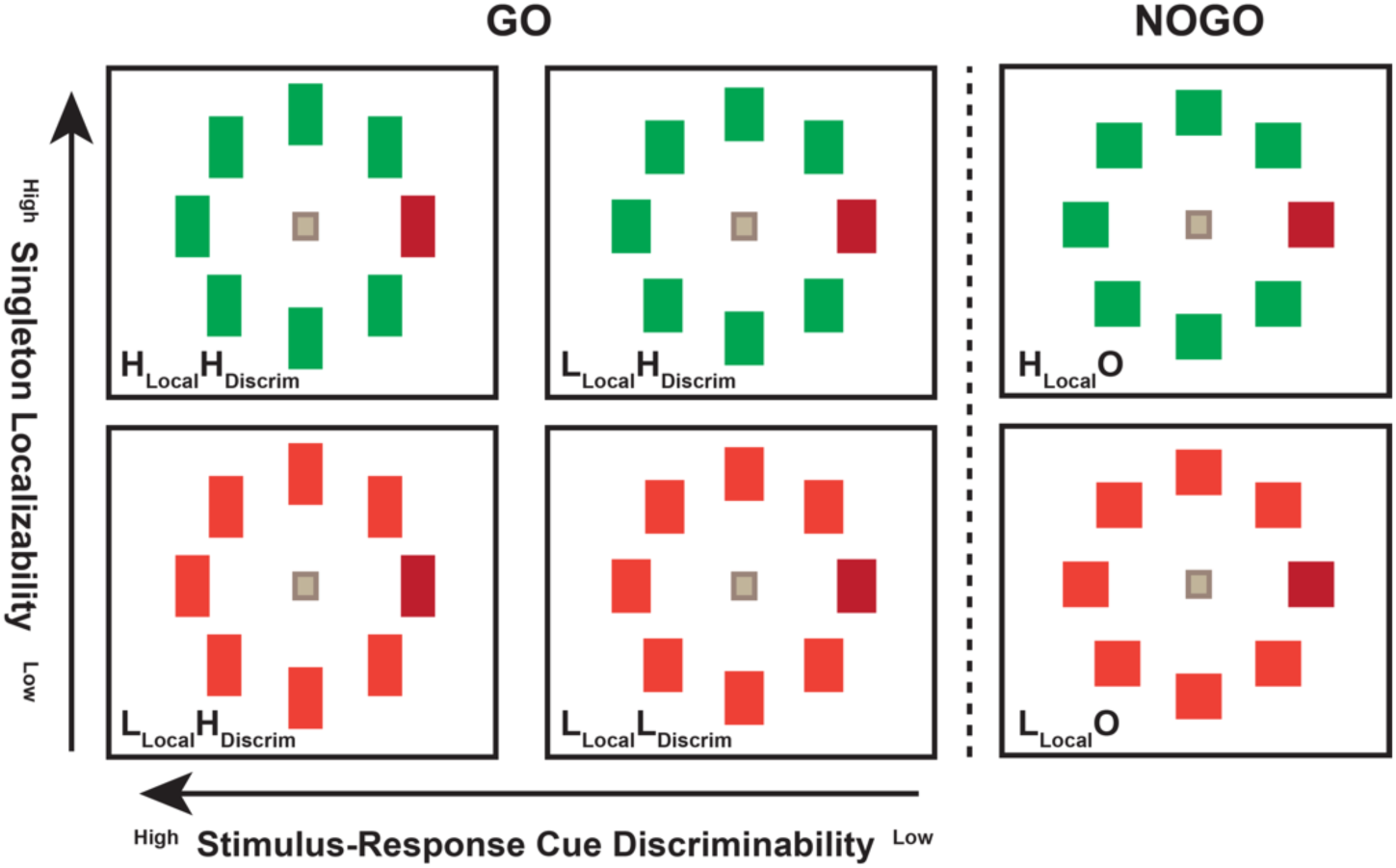
Visual search with separately modified singleton localization and stimulus-response cue discrimination. A) Six basic trial types with singleton illustrated as red. Stimulus elongation cued the response rule – square NOGO, elongated GO (horizontal arrow).

**Table 1.** All possible display combinations with the corresponding difficulty level.

| Color |  | Shape | Difficulty |  |
| --- | --- | --- | --- | --- |
| Singleton | Distractor | All Items | Singleton Localizability | Stimulus-Response Cue Discriminability |
| red | off-red | square | High | NOGO |
|  |  | short rectangle | High | High |
|  |  | long rectangle | High | Low |
|  | green | square | Low | NOGO |
|  |  | short rectangle | Low | High |
|  |  | long rectangle | Low | Low |
|  | off-green | square | Low | NOGO |
|  |  | short rectangle | Low | High |
|  |  | long rectangle | Low | Low |
| off-red | red | square | High | NOGO |
|  |  | short rectangle | High | High |
|  |  | long rectangle | High | Low |
|  | green | square | Low | NOGO |
|  |  | short rectangle | Low | High |
|  |  | long rectangle | Low | Low |
| green | red | square | Low | NOGO |
|  |  | short rectangle | Low | High |
|  |  | long rectangle | Low | Low |
|  | off-red | square | Low | NOGO |
|  |  | short rectangle | Low | High |
|  |  | long rectangle | Low | Low |
|  | off-green | square | High | NOGO |
|  |  | short rectangle | High | High |
|  |  | long rectangle | High | Low |
| off-green | red | square | Low | NOGO |
|  |  | short rectangle | Low | High |
|  |  | long rectangle | Low | Low |
|  | green | square | High | NOGO |
|  |  | short rectangle | High | High |
|  |  | long rectangle | High | Low |

### Data analysis

Gaze position data was filtered with a 2^nd^ order Butterworth low-pass filter with a cut-off frequency of 80 Hz. A velocity threshold of 80°/s was used to identify all saccades. Saccade initiation was the defined as the time when monotonic change in gaze movement began before the velocity reached the threshold. Saccade termination was the time after eye velocity dropped below the threshold and the monotonic change in eye position ended. To be included in analyses, eye movements must have met the following criteria: (1) saccade initiation 80 to 1000 ms after array presentation, (2) peak velocity between 80 and 1000°/s, (3) displacement between 1° and 13° from the fixation point, (4) initial radial position less than 2° from the fixation point.

Saccade vigor was quantified by the relationship between saccade amplitude and peak velocity of all task-relevant saccades. This relationship is typically linear for saccades of displacement up to ∼20° (Bahill et al., 1975; Collewijn et al., 1988). The majority of saccades recorded from both monkeys shifted gaze less than 12°. Therefore, we fitted to the relationship between amplitude and peak velocity a linear function constrained to pass through the origin. We computed the vigor of each saccade as the ratio between the measured velocity and that expected according to the linear fit based on the amplitude (see Reppert et al., 2018). Ratios > 1.0 measure saccades with greater than expected vigor.

We use a chi-square test to assess whether localization error rate and response selection error rate differ significantly across task conditions defined by localizability and cue discriminability. A Least Significant Difference test was used to compare the mean RT between correct and error trials for each condition. A t-test was used to compare the mean vigor between overall correct and localization error saccades, between correct and response selection error saccades, and between localization error response selection error saccades. Two-way ANOVA tests were used to assess the effects of manipulating localizability and discriminability on RT and saccade vigor.

## RESULTS

These results are based on performance measured in 102 sessions (Da: 62 sessions, Le: 40 sessions). One session from Le was removed due to poor gaze data quality, resulting in 101 sessions for subsequent analyses. 1.53% of trials were removed due to artifacts preventing reliable saccade detection.

### Correct response times

The correct RT of both monkeys was separately modified by the localizability and discriminability manipulations (Figure 2; Table 2). RT was significantly longer on trials with higher singleton-distractor similarity and lower response cue discriminability. No significant interaction between singleton localizability and cue discriminability was observed for either monkey. The variation of RT across conditions observed here replicated that reported for 60 earlier sessions (Da 30, Le 30) (Lowe et al., 2019).

**Figure 2.**
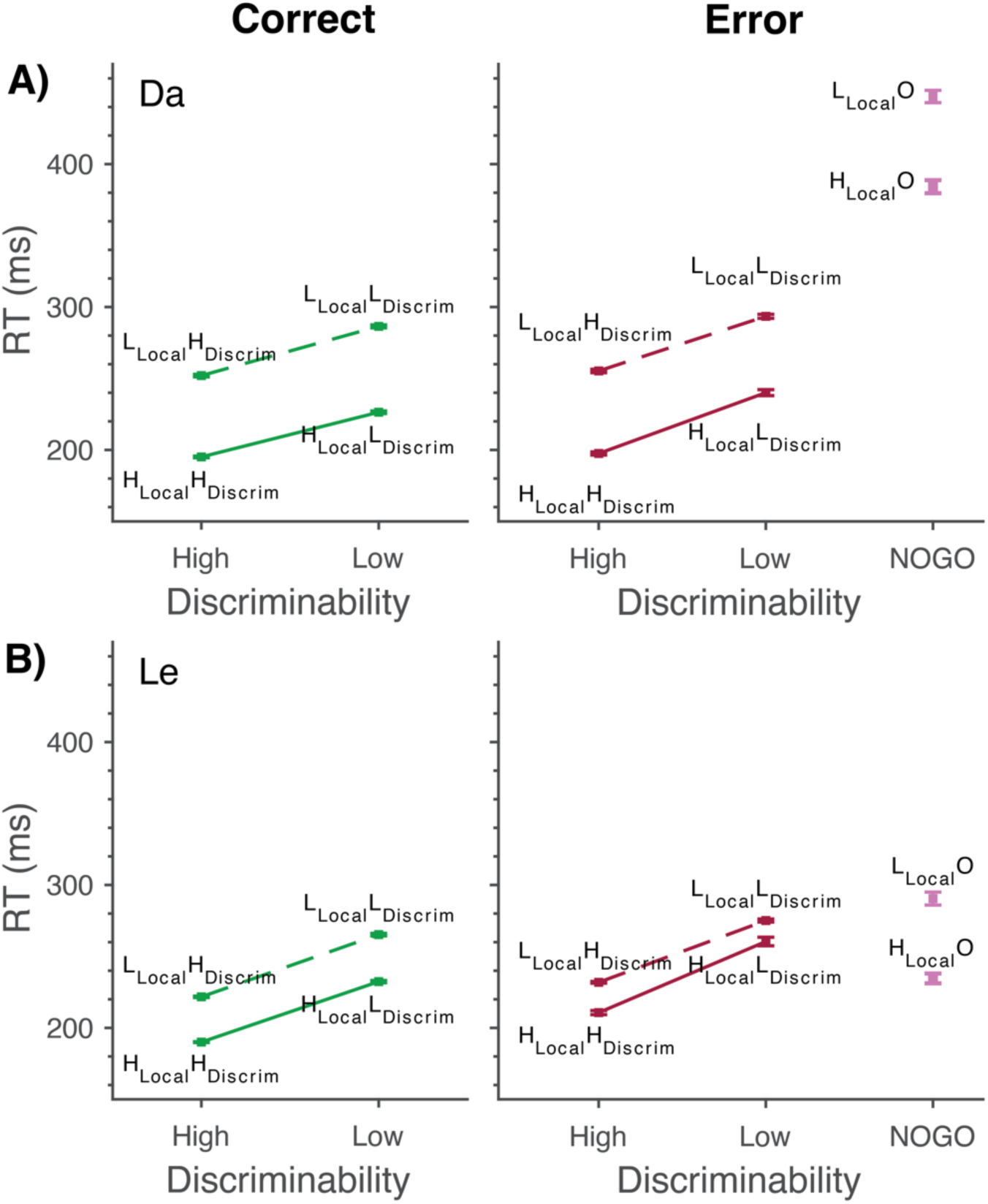
Mean ± SEM RT for correct (Left) and error (Right) responses for Da (A) and Le (B). Correct (green) and Localization Error (red) trials are shown across conditions of varying singleton localizability and cue discriminability: most efficient (H**_Local_**H**_Discrim_**), least efficient (L**_Local_**L**_Discrim_**), and the intermediate conditions (L**_Local_**H**_Discrim_**, H**_Local_**L**_Discrim_**). Response selection errors across conditions of varying singleton localizability (L**_Local_**O, H**_Local_**O) are plotted in magenta. Both localizability and discriminability modulate RT for all trial outcomes.

**Table 2.**
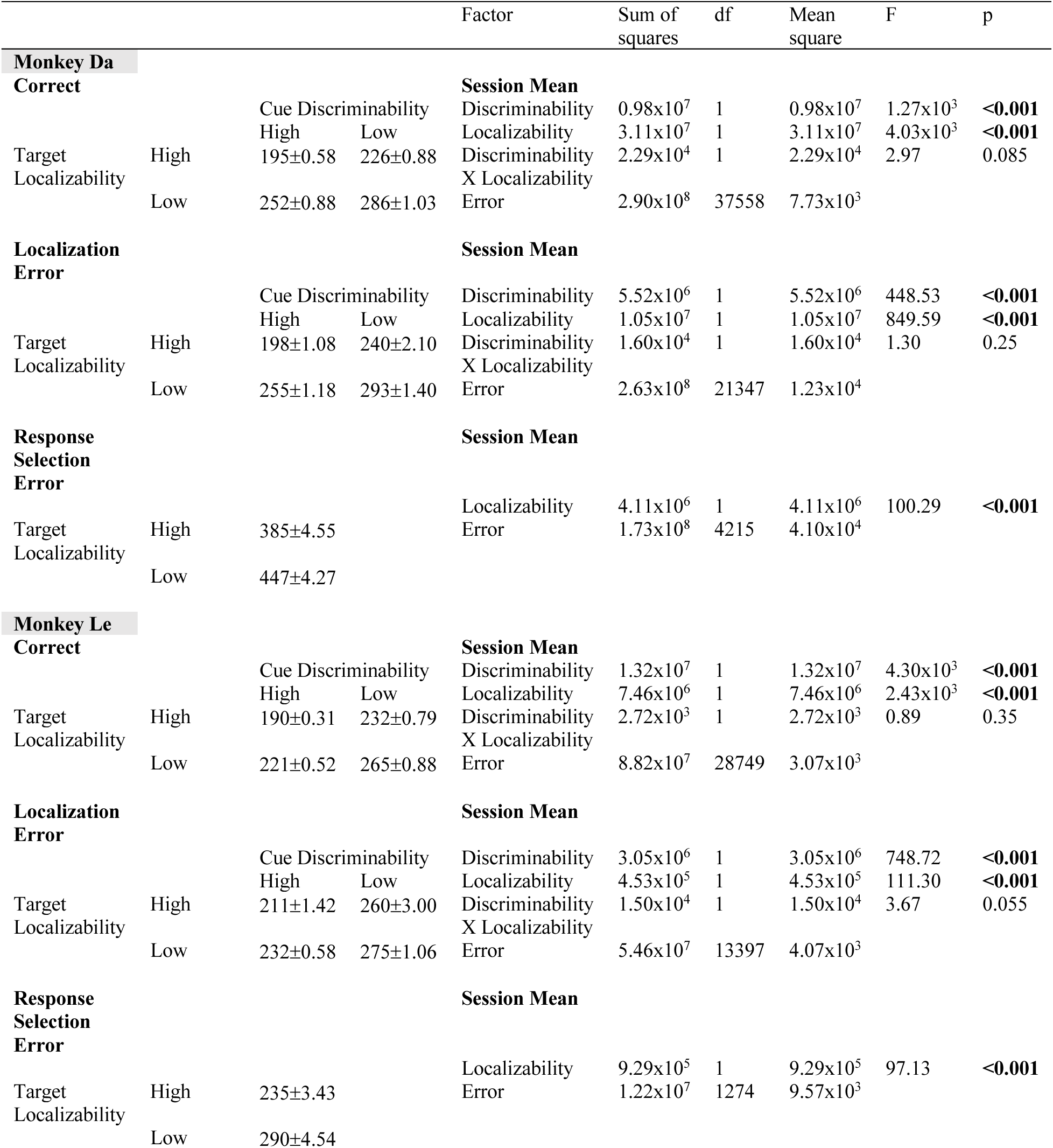
Response time mean ± SEM (ms) and associated ANOVA table.

### Error rates

Overall error rate was 28.3% (Da 29.7%, Le 26.1%). Monkeys produced two errors most commonly, and their incidence varied across the 2x2 factors (Table 3). Both monkeys exhibited failures in the localization process revealed by gaze shifts to a distractor instead of the singleton (Da 36.1% of GO trials, Le 31.8%). Predictably, these errors were more frequent when singleton localizability was low in L**_Local_**H**_Discrim_** and L**_Local_**L**_Discrim_** compared to H**_Local_**H**_Discrim_** and H**_Local_**L**_Discrim_** conditions (Table 3). A chi-square test showed a significant main effect of localizability on localization error rate. The incidence of localization errors was less influenced by cue discriminability. For monkey Da, for both levels of localizability, localization error rates were more frequent when stimulus-response discriminability was more efficient. For Monkey Le, localization error rate was influenced by cue discriminability only when singleton localizability was inefficient, and high discriminability produced more localization errors.

**Table 3.**
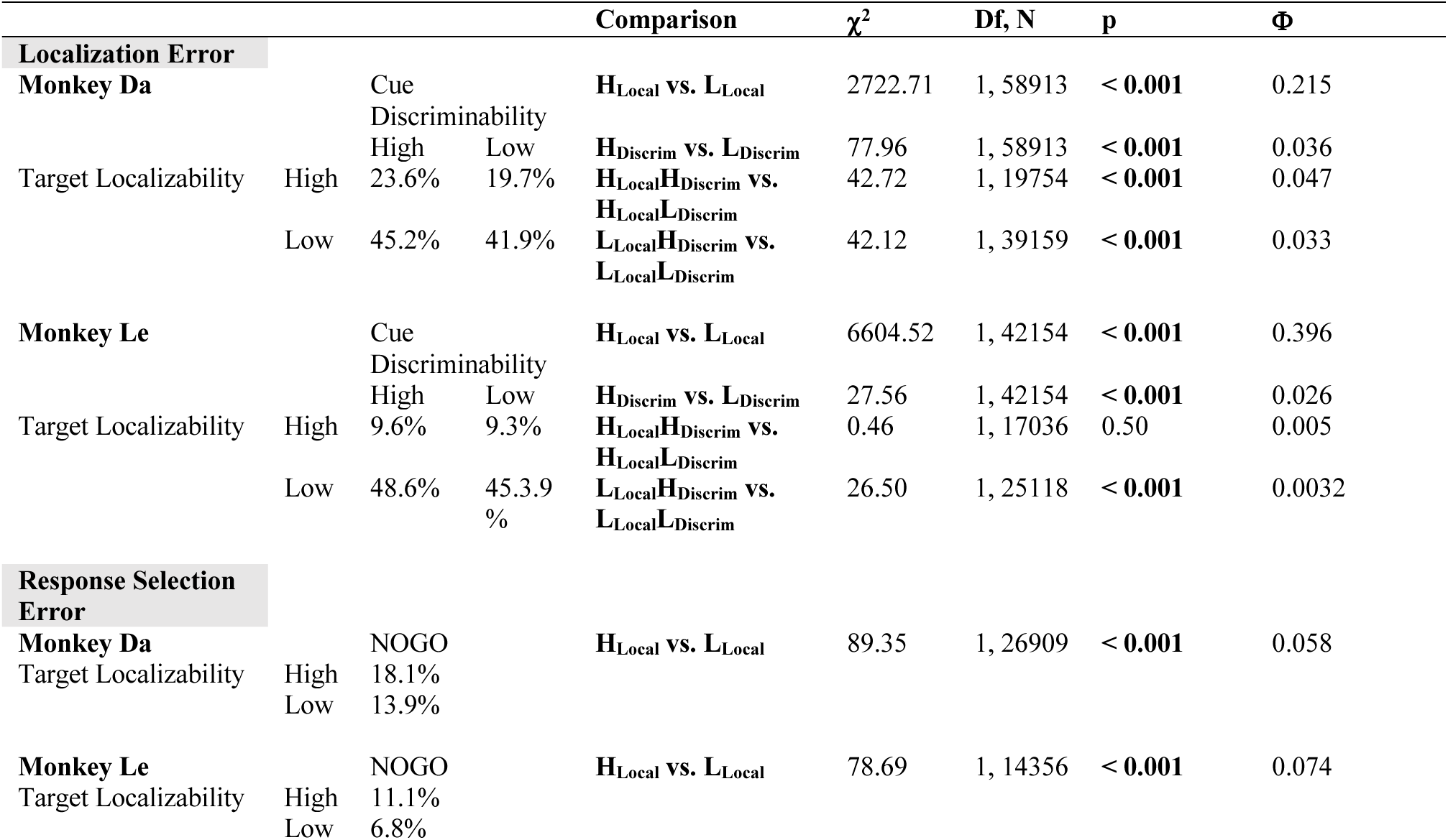
Error rates for localization error and response selection error.

Response selection errors were failures to maintain fixation on NOGO trials. They were more common in Da (15.6% of NOGO trials) than Le (9.0%). Both monkeys made more response selection errors when localizability efficiency was high (Table 3).

### Error response times

For both monkeys, the variation of RT of localization errors across the localizability and discriminability factors mirrored that observed on correct trials with separate modifiability by both the localizability and discriminability manipulations (Figure 2; Table 2). No significant interaction between the two factors was observed. For monkey Da, relative to correct RTs, localization error RTs were significantly longer in the H**_Local_**L**_Discrim_** (mean difference = 13 ms, 95% CI [9, 19], *p* < .001), L**_Local_**H**_Discrim_** (mean difference = 3 ms, 95% CI [1, 6], *p* = .017), and L**_Local_**L**_Discrim_** (mean difference = 7 ms, 95% CI [4, 10], *p* < .001) conditions, but not in the H**_Local_**H**_Discrim_** (*p* = .29) condition. For Le, localization error RTs were significantly longer than correct RTs in all conditions (H**_Local_**H**_Discrim_**: mean difference = 21 ms, 95% CI [16, 25], *p* < .001; H**_Local_**L**_Discrim_**: mean difference = 28 ms, 95% CI [24, 32], *p* < .001; L**_Local_**H**_Discrim_**: mean difference = 10 ms, 95% CI [8, 12], *p* < .001; L**_Local_**L**_Discrim_**: mean difference = 10 ms, 95% CI [8, 12], *p* < .001).

Thus, the RTs of localization error were consistently longer than those of correct RTs. The magnitude of the difference was relatively small in Da, ranging from 1.2 to 6.2%. It was larger in Le, ranging from 3.8 to 12.1%. When cue discriminability was low and target localizability was high, both monkeys produced localization errors with disproportionately longer RTs. This observation indicates that some localization errors arose from strategic rather than sensory processes.

Response selection error RTs varied significantly with singleton localizability (Figure 2; Table 2). For Da but not Le, response selection error RTs were nearly twice the duration of those observed on correct and localization error trials, under both high and low localizability.

### Saccade Vigor

To calculate saccade vigor, we fitted the equation g(x) = α•x to the saccade main sequence across all task-relevant saccades for each monkey across all sessions, producing a monkey-specific mean slop of α [60.30 ± 0.03 (Da) and 69.45 ± 0.05 (Le); mean ± 95% confidence interval] with high goodness of fit: R^2^ = 0.89 (Da) and R^2^ = 0.85 (Le). This one-parameter model accounted for, on average, 87% of the variance in monkeys’ saccade peak velocity. We then computed the ratio of the peak velocity of each saccade to the predicted velocity under the one-parameter model. We also computed saccade vigor using an alternative approached described in Quaia et al. (2000) where variations in peak velocity for saccades directed to the same array eccentricity were compensated in the regression analysis. Computing vigor using this alternative method did not change the main findings.

Figure 3 shows the main sequence of saccade velocity as a function of amplitude with the linear regression for all saccades from one session of monkey Da. A histogram of the distribution of saccade amplitudes is included. The two clusters of saccades were produced to search arrays with eccentricities of 6° and 8°. For monkey Da, the saccade amplitude distribution had a bimodal shape with a left skew. These lower amplitude saccades at the left tail of the distribution primarily came from the response selection error trials. In contrast, monkey Le rarely made low amplitude saccades.

**Figure 3.**
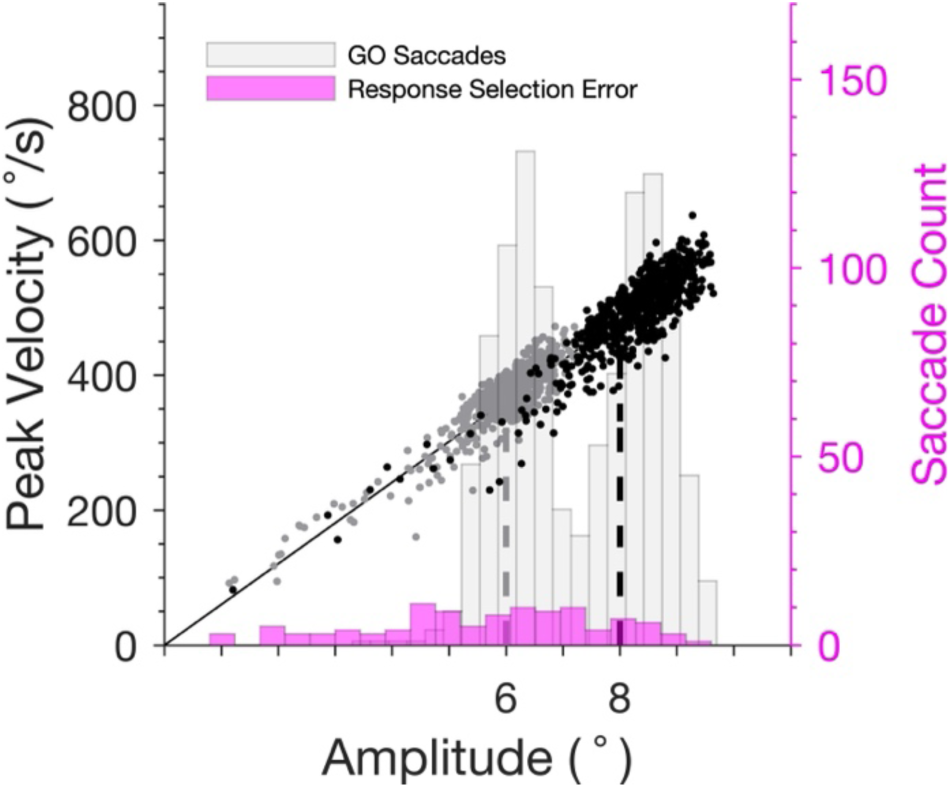
Saccade main sequence from one session from Monkey Da. Saccades were made to a singleton with either 6° (gray) and 8° (black) eccentricity. Left ordinate plots peak saccade velocity. Right ordinate plots the count of saccades across amplitude separately for response selection error saccades (magenta) and for all GO trial saccades, both correct and error (light gray). Shorter saccades at the left tail of the distribution were saccades made on response selection error trials.

Across monkeys, correct saccades exhibited significantly higher average vigor than did localization errors saccades and response selection errors (Table 4). Localization error saccade vigor was also higher than that of response selection error. This variation of vigor across trial outcomes relates to the pattern of RTs. Thus, correct trials showed the fastest mean RT and the highest mean vigor. Localization errors had the second-fastest mean RT and second-highest vigor. Response selection error trials had the slowest mean RT and lowest vigor.

**Table 4.** Vigor mean ± SEM and the t-test comparisons across outcomes.

|  | n | Mean Vigor | T-test Comparison | t | df | p | Cohen's d |
| --- | --- | --- | --- | --- | --- | --- | --- |
| **Monkey Da** |  |  |  |  |  |  |  |
| **Correct** | 38203 | 1.01±<br>3.14x10<sup>-4</sup> | **Correct vs. Localization**<br>**Error** | 15.11 | 60137 | **<0.001** | 0.13 |
| **Localization**<br>**Error** | 21936 | 1.00±<br>4.80x10<sup>-4</sup> | **Correct vs. Response**<br>**Selection Error** | 18.60 | 42418 | **<0.001** | 0.30 |
| **Response**<br>**Selection Error** | 4217 | 0.99±<br>1.68x10<sup>-3</sup> | **Localizability Error vs.**<br>**Response Selection**<br>**Error** | 9.17 | 26151 | **<0.001** | 0.15 |
| **Monkey Le** |  |  |  |  |  |  |  |
| **Correct** | 28753 | 1.01±<br>3.87x10<sup>-4</sup> | **Correct vs. Localization**<br>**Error** | 4.96 | 42152 | **<0.001** | 0.05 |
| **Localization**<br>**Error** | 13401 | 1.01±<br>5.42x10<sup>-4</sup> | **Correct vs. Response**<br>**Selection Error** | 16.09 | 30027 | **<0.001** | 0.46 |
| **Response**<br>**Selection Error** | 1276 | 0.98±<br>2.64x10<sup>-3</sup> | **Localizability Error vs.**<br>**Response Selection**<br>**Error** | 14.24 | 14675 | **<0.001** | 0.42 |

**Table 5.**
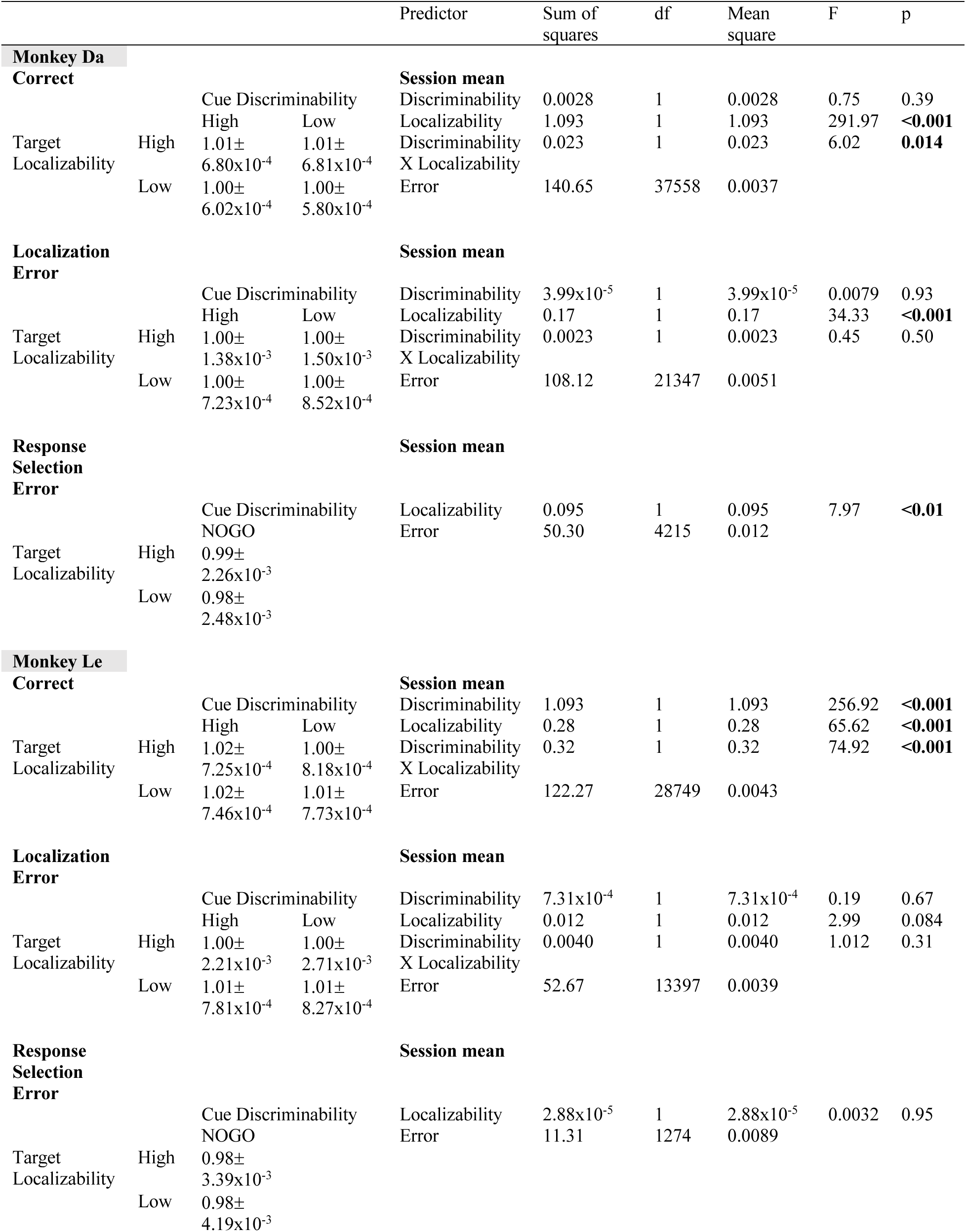
Saccade vigor mean ± SD (ms) and associated ANOVA table.

To relate saccade vigor to RT for each trial outcome, we grouped saccades into 10 RT quantiles and calculated the mean and standard error of RT and vigor for each quantile (Figure 4). Across both monkeys and trial outcomes, saccade vigor decreased with RT. For monkey Da, the overall relationship between vigor and RT was consistent across the correct and error trials. Note that the fastest response selection errors saccades were ∼50 ms slower than the fastest correct and localization error saccades. The absence of the very fast saccades (and typically high vigor) explains why response selection error trials had the lowest mean vigor (Fig. 5A). For monkey Le, the response selection error function significantly deviates from the correct and localization error functions. Saccade vigor was lowest on the response selection error trials across all RTs.

**Figure 4.**
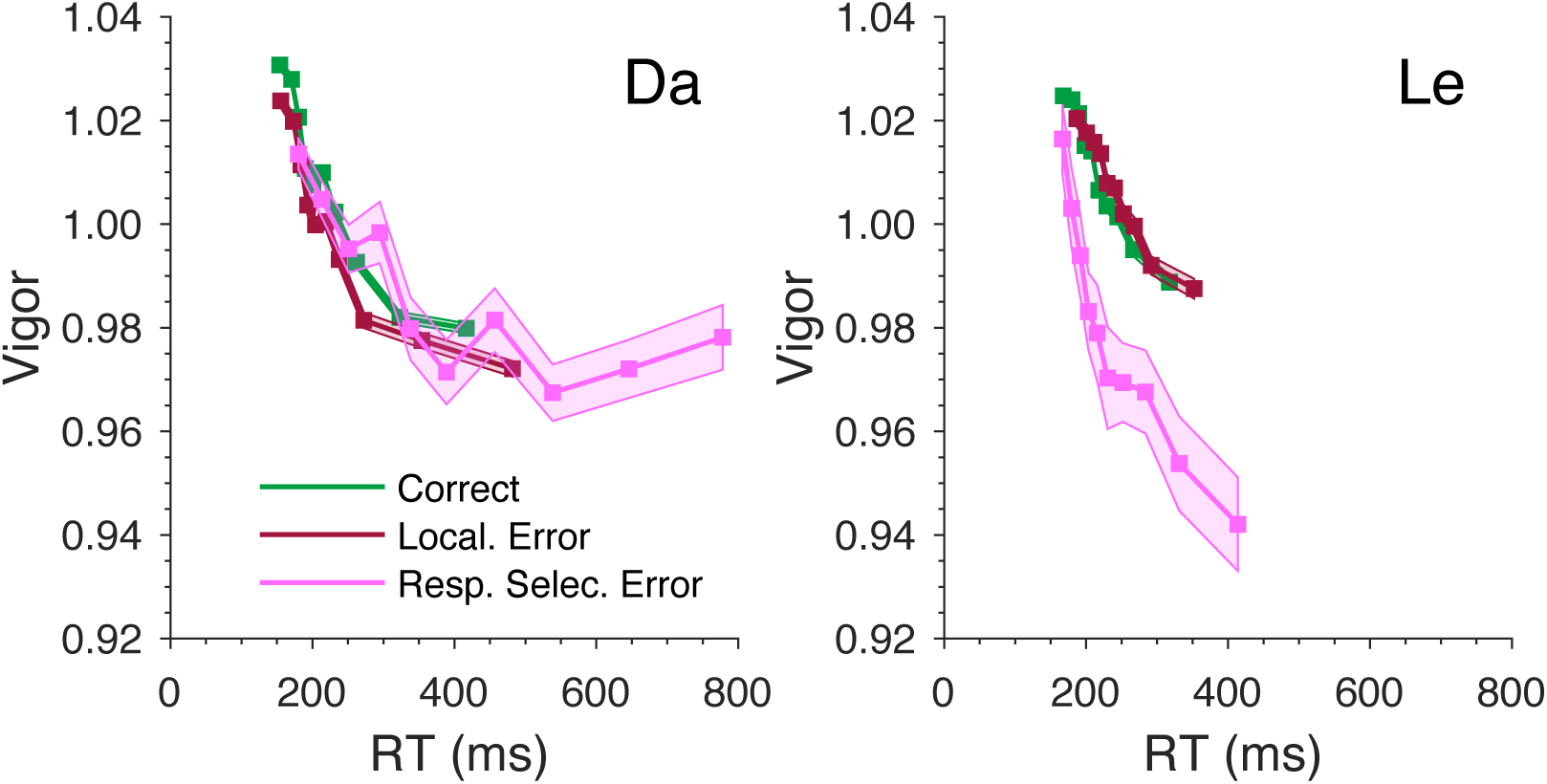
Saccade vigor as a function of RT for Da (left) and Le (right). For each trial outcome, saccades were binned into 10 equally sized bins based on RT percentiles. The mean RT and vigor for each bin as well as the SEM were plotted.

**Figure 5.**
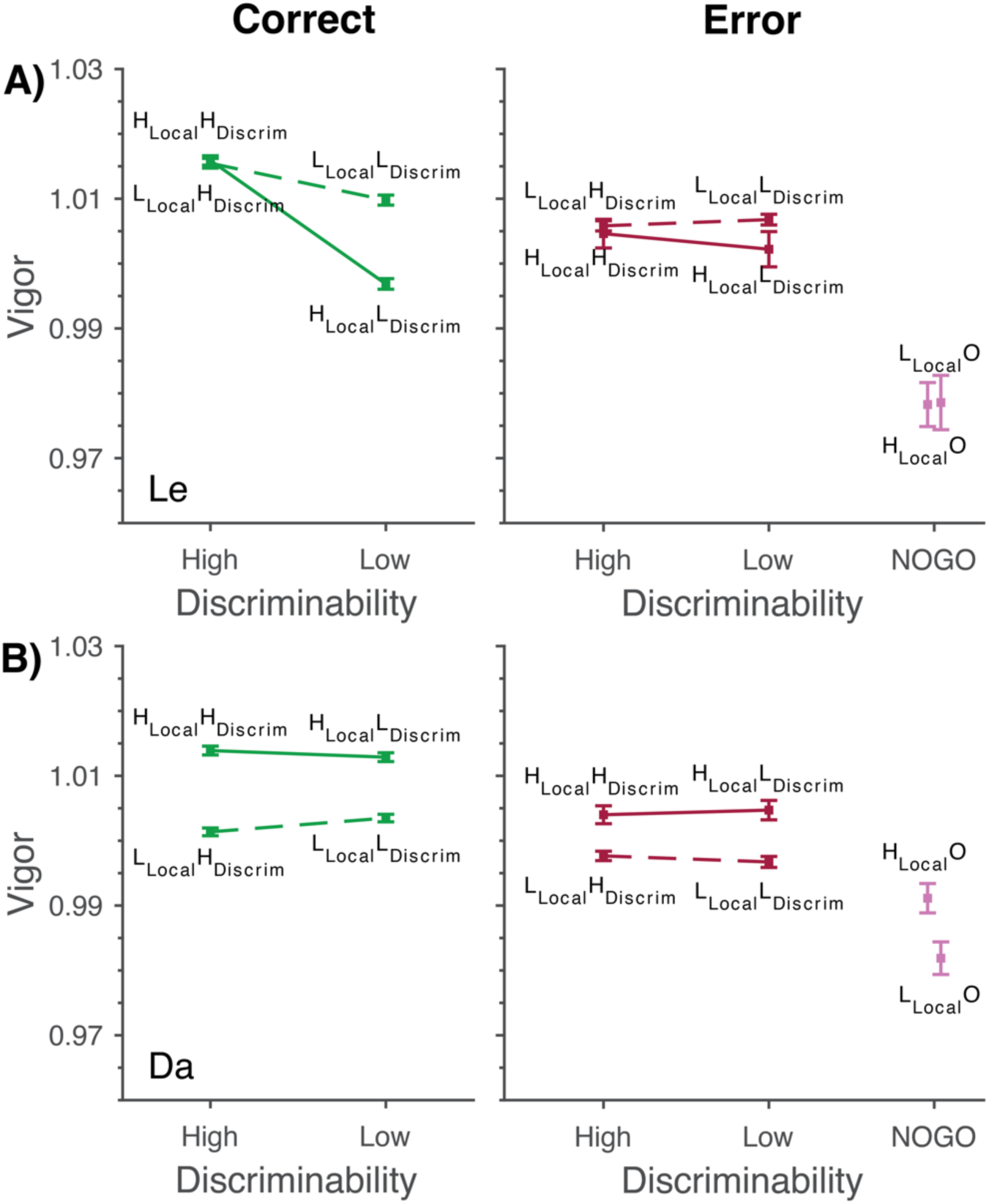
Mean ± SEM Vigor for correct (Left) and error (Right) responses for Da (A) and Le (B). Correct (green) and Localization Error (red) trials are shown across conditions of varying singleton localizability and cue discriminability: most efficient (H**_Local_**H**_Discrim_**), least efficient (L**_Local_**L**_Discrim_**), and the intermediate conditions (L**_Local_**H**_Discrim_**, H**_Local_**L**_Discrim_**). Response selection errors across conditions of varying singleton localizability (L**_Local_**O, H**_Local_**O) are plotted in magenta.

### Testing Separate Modifiability of Saccade Vigor

We assessed saccade vigor across the four difficulty conditions, separately for correct and error trials. Singleton localizability and response cue discriminability affected saccade vigor differently for the two monkeys (Figure 5; Table 4). For monkey Da, on correct trials, vigor varied significantly with localizability but not with discriminability and showed a significant interaction between localizability and discriminability (Figure 5A). On localization and response selection error trials, vigor varied significantly with localizability but not with discriminability and showed no interaction between localizability and discriminability. For monkey Le, on correct trials, vigor varied significantly with localizability and with discriminability, with discriminability showing the stronger effect (*F*_1,28749_ = 256.92, *p* < .001) (Figure 5B). The variation of vigor showed a significant interaction between localizability and discriminability; when shape discrimination was less efficient (L_D_), vigor was higher when the efficiency of localizability was low. However, on localization and response selection error trials, vigor was invariant across conditions. To summarize, localizability consistently influenced saccade vigor for monkey Da across correct and error trials, whereas discriminability had no observable effect. But, for monkey Le, we observed a greater effect of discriminability on saccade vigor when localizability was high relative to low. Such effect was significant only on correct trials.

To illustrate the effects of trial outcome and task manipulation on vigor, we plotted the mean trajectory of saccades made on correct, localization error, and response selection error trials (Figure 6). For monkey Da, we additionally split trials by high versus low localizability, given a main effect of localizability on vigor across all trial outcomes. Correct saccades exhibited the highest velocities. Localization error saccades had a slightly slower velocities, while response selection error saccades were slowest. Monkey Da tended to undershoot the saccade when making response selection errors, resulting in smaller amplitude response selection error saccades on average. For Monkey Le, saccade trajectories across high localizability conditions were plotted given a strong effect of discriminability when localizability was high on correct trials. Correct saccades in the H**_Local_**H**_Discrim_** condition had higher velocities than those in the correct H**_Local_**L**_Discrim_** condition. Localization error saccades in the H**_Local_**H**_Discrim_** condition had slightly higher velocities than did localization errors in the H**_Local_**L**_Discrim_** condition. Response selection errors in the H**_Local_**O condition had the slowest velocities.

**Figure 6.**
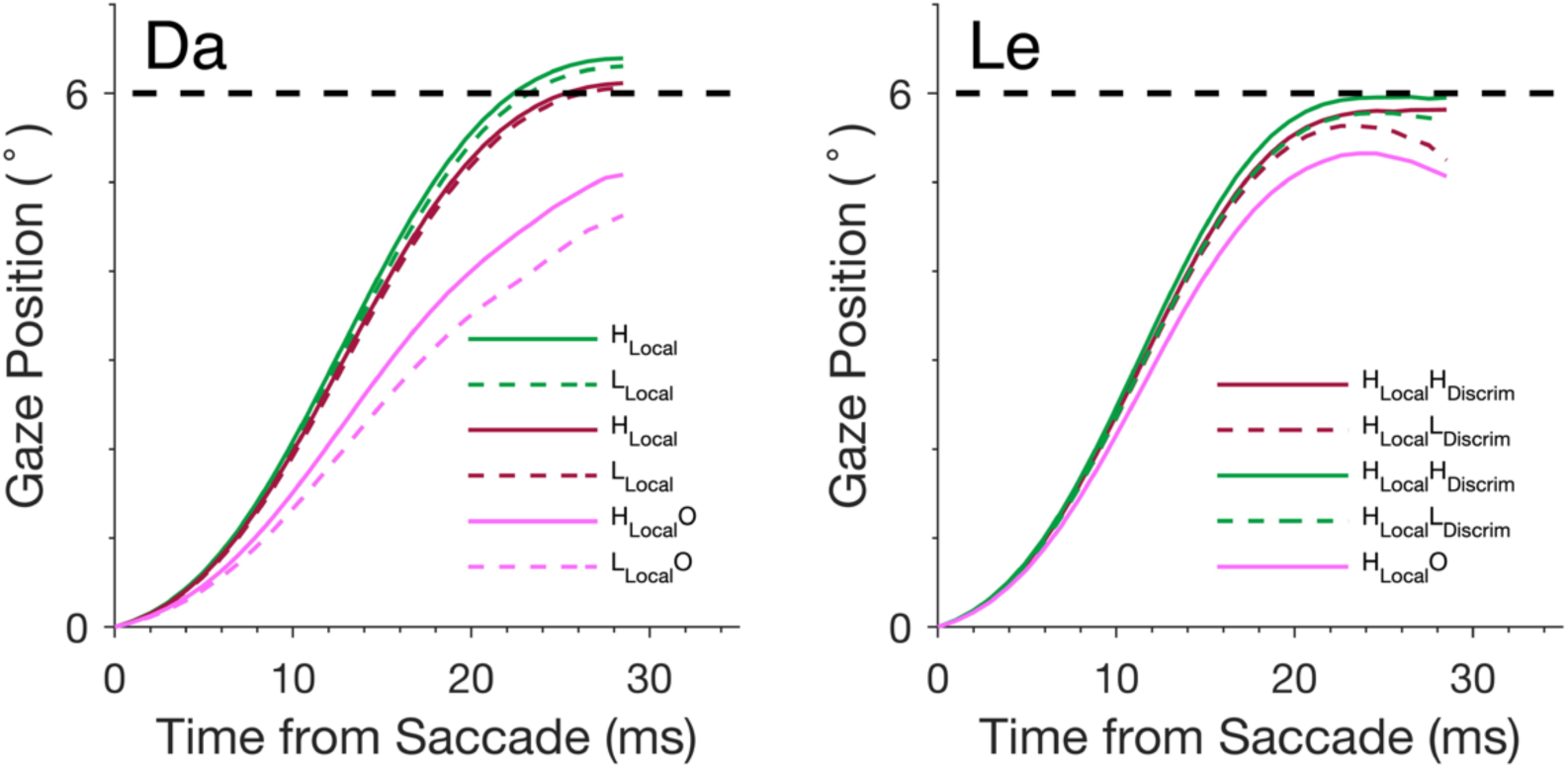
Mean saccade trajectories aligned on saccade onset across selected conditions for Da (left) and Le (right). Saccades are categorized into correct (green), localization error (red), and response selection error (magenta) trials. Only saccades to search array with 6° eccentricity are shown.

## Discussion

Stochastic accumulator models of decision-making treat decision and action as independent, serial processes, but growing evidence shows that response kinematics and dynamics can be influenced by preceding evidence accumulation processes (Dendauw et al., 2024; Molano-Mazón et al., 2024; Reppert, 2025; Seideman et al., 2018; Servant et al., 2021; Wispinski et al., 2020). Using a double-factorial visual search task (Lowe et al., 2019), we show that when multiple evidence accumulation processes guide gaze, the efficiency of each can contribute to the vigor of the gaze shift.

Evidence from non-ballistic movements (e.g., rat body orienting movements and human limb movements) has demonstrated that motor output is continuously shaped by ongoing evidence accumulation (Molano-Mazón et al., 2024). We extended these findings to ballistic eye movements. Our findings are consistent with previous studies demonstrating that saccade vigor is influenced by decision-related variables such as confidence (Seideman et al., 2018), urgency (Thura, 2020), subjective reward (Reppert et al., 2015), effort (Shadmehr et al., 2019), and speed-accuracy trade-off (Reppert, Servant, et al., 2018). Stronger signal strength with longer processing time is associated with more vigorous saccades (Molano-Mazón et al., 2024; Seideman et al., 2018) even if they are not relevant for the task (Joo et al., 2016; Matsumiya & Furukawa, 2023).

Coupling between decision speed and movement vigor has been cited as evidence for the co-regulation of decision and action processes (Fievez et al., 2024; Reppert, 2025). Sometimes, decision and movement speed can also compensate each other (Saleri & Thura, 2024). The presence or absence of co-regulation is contingent on task incentives (Fievez et al., 2024). Both monkeys exhibited the most vigorous saccades on trials with the fastest RT, and mean vigor decreased as RT increased. This inverse relationship was observed on correct and both types of error trials. A prior study of visual search with instructed speed-accuracy trade-off found that saccade vigor varied with RT under the fast but not the accurate instruction (Reppert et al. 2018; see also Carsten et al., 2023). Hence, one interpretation of the present results is that the two monkeys performed this task with an emphasis of speed over accuracy. This interpretation is consistent with their relatively high error rates.

While we observed an overall co-regulation of decision and movement speed in both monkeys, factorial manipulation of the two decision processes affected RT and saccade vigor differently. Manipulating the efficiency of target localization and of response cue discrimination process had an overall additive effect on mean RT for both monkeys. However, only localizability but not discriminability significantly influenced saccade vigor for Da, and such effect was observed across correct, localization error, and response selection error trials. In contrast, monkey Le showed a greater effect of discriminability on vigor when localizability was high relative to low, and only on correct trials. This suggests that while RT and vigor can be influenced by shared underlying mechanisms, the processes of generating decision and movement are distinct and can be modulated independently. Distinct patterns of error rates and saccade vigor across monkeys indicated individual differences in decision-making strategies, which can evolve with training (see Lowe et al., 2019). Individual variability across monkeys have been observed in other decision-making task (e.g., Saleri & Thura, 2024), and previous research has described individual differences in movement vigor generally (Reppert, Rigas, et al., 2018).

Stochastic accumulator models are the current dominant theoretical framework of perceptual decision-making. They treat motor responses as merely the output of the preceding evidence accumulation process reaching a decision threshold. Canonical forms of these models incorporate no interaction between decision variables and response production parameters. Afferent and efferent stages before and after the evidence accumulation process are reduced to a single non-decisional residual component (Ratcliff, 1978, 2013; Ratcliff et al., 2016). However, studies suggest that non-decision time parameters are unlikely to be only sensorimotor delays (Bompas et al., 2024; Smith & Lilburn, 2020; Weindel et al., 2021). More generally, traditional models of decision-making based on discrete RT and accuracy responses (Forstmann et al., 2016; McClelland, 1979; Ratcliff, 1978; Ratcliff et al., 2016) cannot capture internal processes occurring concurrently and continuously (Cisek, 2006; Gallivan et al., 2018; Song & Nakayama, 2009; also Schall, 2019).

Other models have been formulated to approach greater neurobiological plausibility. For example, the interaction between neurons representing visual salience and neurons preparing to initiate saccades has been articulated by a gated accumulator model (Purcell et al., 2010, 2012). This model demonstrates how the neurons preparing saccades correspond to evidence accumulators that trigger saccade initiation (see also Middlebrooks et al., 2020; Ratcliff et al., 2007). This functional attribution is consistent with anatomical observations that these pre-movement evidence accumulation neurons, which are found in structures like the frontal eye field and superior colliculus, send axons to terminate in brainstem circuits responsible for generating saccades. The understanding of the brainstem saccade circuit is intricate and precise (e.g., Daye et al., 2013; Quaia et al., 1999). However, we do not know how the reaching of a threshold by pre-movement neurons causes the initiation and production of saccades (Schall & Paré, 2021). The current findings demonstrate that this linkage is leaky and offer data that would usefully constrain a new integrative model that formally links the evidence accumulator with the brainstem saccade circuit.

Other recent studies and models have explored how decision-making processes can interact with the sensorimotor system to produce systematic covariation in RT and execution (Gallivan et al., 2018; Wispinski et al., 2020). This covariation can be explained by models that incorporate dynamics of both evidence accumulation and responses execution (Dendauw et al., 2024; Molano-Mazón et al., 2024; Servant et al., 2021). If decision and motor output are not strictly sequential processes, then mathematical models must also predict the rate of muscular contraction and explain the kinematics of movement production in relation to decision processes.

To conclude, this study demonstrates that manipulating the quality of decision evidence used for a visual search task influenced both the decision time as well as the saccade dynamics. We separately manipulated the efficiency of two concurrent decision processes—target localization and response cue discrimination. These factors affected RT and saccade vigor differently, indicating the processes underlying response selection and movement execution are distinct and can be regulated somewhat independently. These findings challenge traditional stochastic accumulator models that assume a sequential and independent relationship between decision and action. Movement vigor offers a novel and valuable constraint on biologically plausible models of decision making, which can help address the persistent challenge of model mimicry.

## ACKNOWLEDGEMENTS

The data were collected by K. Lowe and T.R. Reppert with funding by NIH grants F32-EY028846, T32-EY007135, P30-EY08126, and R01-EY00890. Preparation of this manuscript was funded by Ontario Graduate Scholarship to W. Lyu, by Natural Sciences and Engineering Research Council of Canada Undergraduate Research Award to H. Parmar, and by Natural Sciences and Engineering Research Council of Canada grant number: RGPIN-2022-04592.

## Conflict of interest

We have no conflicts.

## Notes

### Competing Interest Statement

The authors have declared no competing interest.

